# VIP-HL: Semi-automated ACMG/AMP variant interpretation platform for genetic hearing loss

**DOI:** 10.1101/2020.08.10.243642

**Authors:** Jiguang Peng, Jiale Xiang, Xiangqian Jin, Junhua Meng, Nana Song, Lisha Chen, Ahmad Abou Tayoun, Zhiyu Peng

## Abstract

**Purpose:** The American College of Medical Genetics and Genomics, and the Association for Molecular Pathology (ACMG/AMP) have proposed a set of evidence-based guidelines to support sequence variant interpretation. The ClinGen hearing loss expert panel (HL-EP) introduced further specifications into the ACMG/AMP framework for genetic hearing loss. This study aimed to semi-automate the HL ACMG/AMP rules.

**Methods:** VIP-HL aggregates information from external databases to automate 13 out of 24 ACMG/AMP rules specified by HL-EP, namely PVS1, PS1, PM1, PM2, PM4, PM5, PP3, BA1, BS1, BS2, BP3, BP4, and BP7.

**Results:** We benchmarked VIP-HL using 50 variants where 83 rules were activated by the HL expert panel. VIP-HL concordantly activated 96% (80/83) rules, significantly higher than that of by InterVar (47%; 39/83). Of 4948 ClinVar star 2+ variants from 142 deafness-related genes, VIP-HL achieved an overall variant interpretation concordance in 88.0% (4353/4948). VIP-HL is available with a user-friendly web interface at http://hearing.genetics.bgi.com/.

**Conclusion:** VIP-HL is an integrated online tool for reliable automated variant classification in hearing loss genes. It assists curators in variant interpretation and provides a platform for users to share classifications with each other.

## Introduction

Hearing loss is a primary global health concern, affecting about 6.8% of the world’s population^1^. Genetic factors account for at least 50 to 60% of childhood hearing loss^2^. To date, over 150 deafness-related genes were discovered.^3^ The advancement of next-generation sequencing technology and its continually decreasing cost have facilitated the genetic diagnosis of hearing loss.^4^ One of the remaining significant challenges, however, is the accurate interpretation of a large number of identified variants.

In 2015, the American College of Medical Genetics and Genomics and the Association for Molecular Pathology (ACMG/AMP) published a joint guideline to standardize variant interpretation.^5^ The guideline offered a common framework for curators to resolve variant classification differences, and to improve interpretation consistency across laboratories.^6^ To evolve the guideline over time, National Institutes of Health-funded Clinical Genome Resource (ClinGen) developed groups of disease experts to refine recommendations for different disease areas, including genetic hearing loss,^7^ *MYH7-associated* inherited cardiomyopathies,^8^ germline *CDH1*,^9^ and *PTEN-associated* hereditary cancer.^10^

Using bioinformatics tools to facilitate and standardize variant interpretation is one of top priorities, and has been shown to be useful for curators.^11^ InterVar is one such tool semi-automating 18 criteria in the ACMG/AMP guidelines.^12^ However, generic tools may not fulfill the need because many interpreting standards are disease-specific and vary dramatically amongst different diseases.^13^ In 2018, two tools named CardioClassifier^14^ and CardioVAI^15^ were explicitly developed for interpreting variants in cardiovascular diseases. In 2019, a semi-automated tool called “Variant Interpretation for Cancer” (VIC) was developed to accelerate the interpretation process in Cancer.^16^

Here, we developed a **V**ariant **I**nterpretation **P**latform for genetic **h**earing **l**oss (VIP-HL). This new online tool utilizes the framework outlined by the ClinGen Hearing Loss Expert Panel (HL-EP)^7^ to automatically annotate variants in 142 hearing loss related genes across 13 ACMG/AMP rules. VIP-HL is freely accessible for non-commercial users in a web server at http://hearing.genetics.bgi.com/.

## Methods

VIP-HL automates 13 out of 24 ACMG/AMP rules specified by ClinGen HL-EP, namely PVS1, PS1, PM1, PM2, PM4, PM5, PP3, BA1, BS1, BS2, BP3, BP4, and BP7 (Oza et al. 2018). The development and optimization of VIP-HL are described in three sections: (1) rule selection, optimization and implementation; (2) Benchmarking and comparative analysis; (3) web interface development.

### Rule selection, optimization and implementation

Twenty-four ACMG/AMP rules for genetic hearing loss were grouped into four sections: population (PM2, BA1, BS1, BS2), computational (PVS1, PS1, PM1, PM4, PM5, PP3, BP3, BP4, and BP7), case/segregation (PM3, PS2/PM6, PS4, PP1, PP4, BS4, BP2, and BP5) and functional (BS3 and PS3) data. VIP-HL automates all 13 population and computational rules, while case/segregation and functional criteria (n=11) still require manual curation.

For population rules (PM2, BA1, and BS1), our implementation of population frequency cutoffs relied on gnomAD v2.1 exomes and genomes combined dataset^17^. The cutoffs followed the guideline for genetic hearing loss ^7^ The highest filtering allele frequency across all gnomAD populations (“popmax”) was applied as the allele frequency for each variant.^18^ To determine the BS2 criterion, we retrieved the homozygote number from the gnomAD control dataset. BS2 was applied when the homozygote number was greater than three for autosomal recessive disease, and one for autosomal dominant disease. It should be noted that ClinGen HL-EP did not determine the cutoffs of the homozygous number for BS2. The default setting by VIP-HL can be adjusted by users.

The parameterization of nine computational criteria (PVS1, PS1, PM1, PM4, PM5, PP3, BP3, BP4, and BP7) was described as follows. The specifications of PVS1 were, clarified before.^19,20^ PM1 can be applied when a variant is in a mutational hotspot region or well-studied functional domain without benign variation.^5,7^ The hotspot region/domain was determined based on the enrichment of pathogenic variants, as clarified in our recent work.^19^

Pathogenic/likely pathogenic variants from ClinVar 20200629 release were retrieved to determine the existence of the same amino acid changes (PS1) and the different amino acid changes (PM5). For example, knowing that NM_004004.6(*GJB2*):c.109G>A (p.Val37Ile) is a well-established pathogenic variant in ClinVar, we can now apply PM5 for NM_004004.6(*GJS2*):c.109G>T (p.Val37Phe).

PM4 was applied when the in-frame deletion or insertion was not in the repetitive region which is evolutionary well conserved (Oza et al. 2018). The repetitive region was determined based on RepeatMasker^21^ and Tandem repeats finder^22^. The region with GERP scores higher than two was considered as evolutionarily conserved.^23^ BP3 was applied while the in-frame indels were in repeat region without known function.

In silico tools selected for implementing PP3, BP4 and BP7 were REVEL^24^ and MaxEntScan^25^. PP3 was applied with a REVEL score of ≥0.7 and BP4 was applied with a REVEL score≤0.15.^7^ PP3 can also be applied when non-canonical splice variants were predicted to have an impact on splicing via MaxEntScan.^25^ BP7 was employed when a synonymous variant was predicted with no impact on splicing via MaxEntScan and the nucleotide is not highly conserved (GERP < 2).^26^

The final pathogenicity is reported in five tiers system proposed by the ACMG/AMP guideline, namely “Pathogenic (P)”, “Likely Pathogenic (LP)”, “Variant of Uncertain Significance (VUS)”, “Likely Benign (LB)”, and “Benign (B)”.^5^

### Benchmarking and comparative analysis

In order to test VIP-HL performance, we constructed two benchmark datasets. The first dataset consisted of 50 out of 51 variants in the hearing loss gene that has been curated by the ClinGen HL-EP.^7^ We excluded NM_206933.3:c.(?_12295)_(14133_?)del in the *USH2A* gene because it is an exon-level deletion (Exons 63-64 deletion), which is currently not compatible with VIP-HL.

To assess the importance of disease-specific annotations, we compared activated rules by ClinGen HL-EP with those activated by VIP-HL and InterVar, and vice versa. Comparing rules that were not activated by ClinGen HL-EP, but activated by either VIP-HL or InterVar, we did not count the variants meriting the BA1 criterion because ClinGen HL-EP did not activate other criteria once a variant met the BA1 criterion. For example, ClinGen HL-EP assigned BA1 for NM_005422.2:c.1111A>G in the *TECTA* gene and did not further activate other criteria. However, both VIP-HL and InterVar activated BS2 because 10518 homozygotes are reported in the gnomAD database for this variant.^17^ The InterVar code was downloaded from GitHub. All the settings were set as default.

The second dataset included 4948 variants in 142 deafness-related genes with ClinVar star 2+ (i.e., multiple submitters with assertion criteria, expert panel or practice guideline).^27^ These variants were selected because they had fewer misclassifications.^28,29^ The 142 deafness-related genes were curated by ClinGen HL-EP.^30^ The gene list and their gene-disease associations are listed in Supplementary Table 1.

### Web interface development

The front end of VIP-HL is written in HTML5 and JavaScript (VUE), with the back end implemented in JAVA and Python 3. Uploaded variants are annotated via Ensembl Variant Effect Predictor.^31^ A total of 142 hearing loss genes as mentioned above were collected from the ClinGen website (DiStefano et al. 2019). We pre-annotated all variants in these genes from ClinVar 20200629 release. For variants not pre-annotated, VIP-HL can execute the algorithm and provide the interpreting results promptly.

## Results

### Benchmark analysis

Adopting ACMG/AMP guidelines for genetic hearing loss, 152 ACMG/AMP rules were activated for 50 pilot variants curated by the ClinGen HL-EP. Of these, 55% (83/152) rules could be automatically interpreted by VIP-HL, spanning PVS1, PM1, PM2, PM4, PM5, PP3, BA1, BS1, BP4, and BP7. Overall, VIP-HL achieved 96% (80/83) concordant interpretations compared with activations by ClinGen HL-EP (Figure 1a). Of 83 activated rules (Figure 1b), the three discordant activations between VIP-HL and ClinGen HL-EP were PM2_Supporting (2 times), and BA1 (1 time). Those discrepancies were most likely due to the adoption of popmax filtering allele frequency (Whiffin et al. 2017) by VIP-HL (Table 1, variant #1-3). By comparison, InterVar only achieved 47% (39/83) concordant activations, significantly lower than that by VIP-HL (Figure 1b).

**Figure 1.**
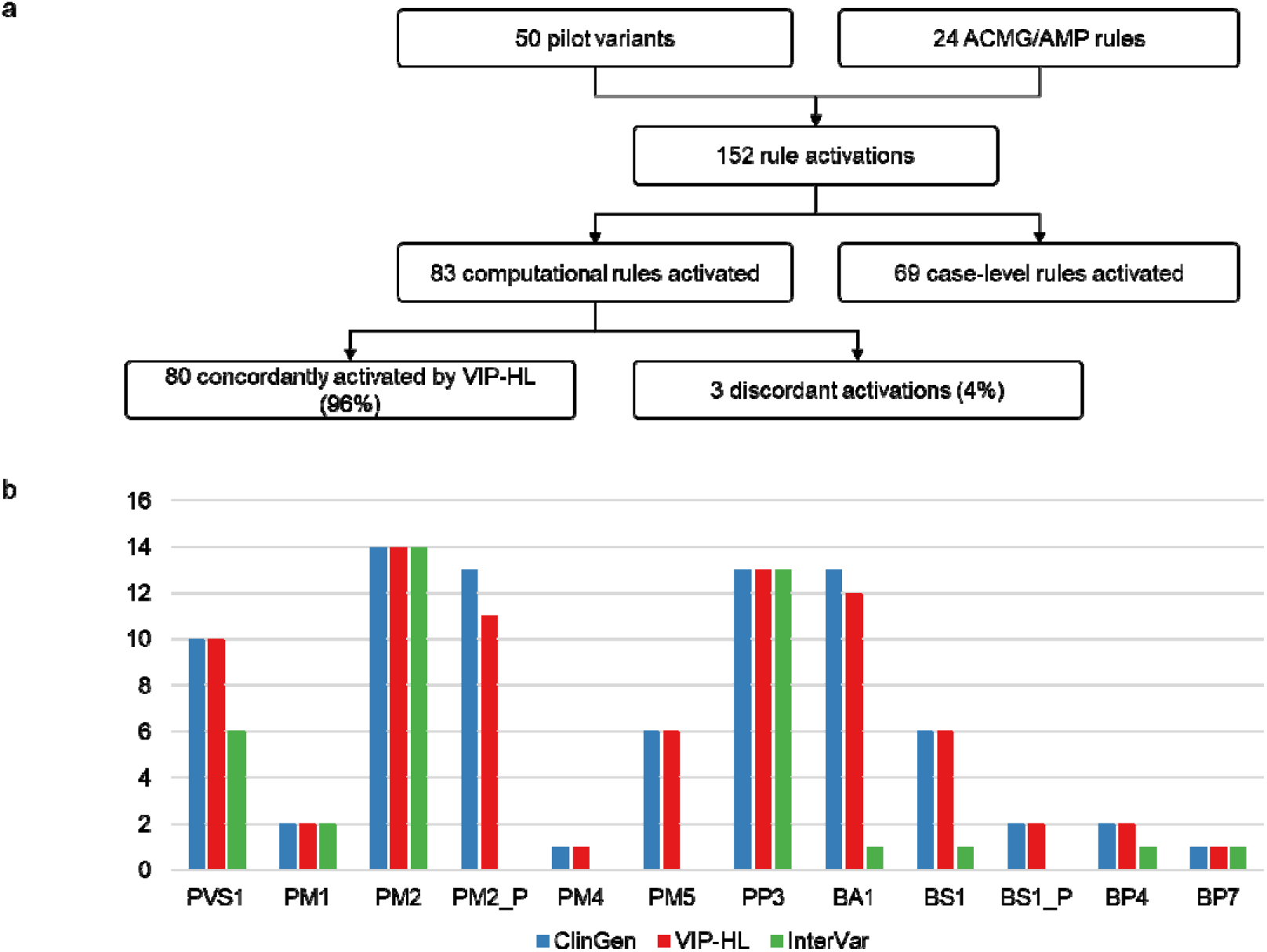
Validation of VIP-HL. (a) Comparing VIP-HL to a set of 50 hearing loss expert panel curated variant. (b) Counts of individual rules activated by ClinGen hearing loss expert panel, VIP-HL and InterVar for 50 pilot variants. Rules applied with a modified strength are denoted by the rule followed by _P for Supporting.

**Table 1.** Analysis of rules activated by VIP-HL and ClinGen Hearing Loss Expert Panel (HL-EP). Four variants (#1-#3) have discrepant rules between ClinGen HL-EP and VIP-HL, spanning BA1 and PM2_P. Five variants (#4-#8) have rules that were not activated by ClinGen HL-EP, but activated by VIP-HL.

| # | Gene | Transcript | c.DNA | Rules activated by ClinGen HL-EP | Rules activated by VIP-HL | Explanations for discordant rules |
| --- | --- | --- | --- | --- | --- | --- |
| 1 | *MYO6* | NM_004999.3 | c.2836C>T | BA1 | BS1 | The popmax filtering allele frequency is 0.00075 in Other in gnomAD. |
| 2 | *USH2A* | NM_206933.2 | c.11241C>A | PVS1, PM2_P | PVS1, PM2 | The popmax filtering allele frequency is 0.000054 in African-American in gnomAD. |
| 3 | *USH2A* | NM_206933.2 | c.14419G>A | PM2P | PM2 | The popmax filtering allele frequency is 0.000064 in South Asian in gnomAD. |
| 4 | *GJB2* | NM_004004.5 | c.101T>C | PM5 | PM5, PM1, PP3, BS2 | PM1 is applied because this variant locates in a mutational hotspot region: 8 pathogenic missense variants and 0 benign missense variant in chr13:20763603-20763633.<br>PP3 is activated because REVEL score is 0.702, greater than the threshold (0.7) that ClinGen Hearing Loss Expert Panel recommends for PP3.<br>BS2 was activated because 16 homozygotes are reported in the gnomAD control dataset. |
| 5 | *GJB2* | NM_004004.5 | c.109G>A | PM5 | PM5, PM1, BS2 | PM1 is applied because this variant locates in a mutational hotspot region: 8 pathogenic missense variants and 0 benign missense variant in chr13:20763603-20763633.<br>BS2 was activated because 50 homozygotes are reported in the gnomAD control dataset. |
| 6 | *GJB2* | NM_004004.5 | c.-22-2A>C | BS1 | BS1, PVS1 | GT-AG 1,2 splice sites -> Exon skipping or use of a cryptic splice site disrupts reading frame and is predicted to undergo NMD -> Exon is present in biologically relevant transcript(s) -> PVS1 |
| 7 | *KCNQ4* | NM_004700.3 | c.720C>G | BP4, BP7 | BP4, BP7, PM2 | PM2 is applied because this variant does not exist in gnomAD. |
| 8 | *KCNQ4* | NM_004700.3 | c.853G>A | PM2, PM5, PM1 | PM2, PM5, PM1, PP3 | PP3 is activated because REVEL score is 0.793, greater than the threshold (0.7) that ClinGen Hearing Loss Expert Panel recommends for PP3. |

Both VIP-HL and InterVar activated certain rules that ClinGen HL-EP did not apply. Specifically, VIP-HL activated only four rules six times, including PM1 (2 times), PP3 (2 times), PM2 (1 time), PVS1 (1 time), BS2 (2 times) (Table 1, variant #4-8). InterVar, however, activated 11 rules 100 times (Supplementary Figure S1), spanning PVS1 (2 times), PM1 (15 times), PM2 (20 times), PP2 (2 times), PP3 (12 times), PP5 (25 times), BS1 (2 times), BS2 (8 times), BP1 (11 times), BP4 (1 times), BP6 (2 times). Notably, four rules (PP2, PP5, BP1, and BP6) that were considered not applicable for genetic hearing loss accounted for 40% of the activations (40/100). These results demonstrated the reliable annotations by VIP-HL and reinforced the importance of disease-specific annotations in variant interpretation.

### Comparison with ClinVar interpretation

Of 4948 ClinVar variants in 142 deafness-related genes, VIP-HL achieved an overall variant interpretation concordance of 88.0% (4353/4948). The concordant interpretations were 57.1% (376/658) in P/LP variants, 93.6% (3083/3295) in B/LB variants, and 89.8% (894/995) in VUS variants, respectively (Table 2).

**Table 2.** Illustration of automated interpretation of variants submitted in ClinVar.

| ClinVar | VIP-HL (automated interpretation) |  |  | All |
| --- | --- | --- | --- | --- |
|  | Pathogenic or Likely pathogenic | Uncertain Significance | Benign/Likely benign |  |
| Pathogenic or Likely pathogenic | 376 | 280 | 2 | 658 |
| Uncertain Significance | 4 | 894 | 97 | 995 |
| Benign/Likely benign | 1 | 211 | 3083 | 3295 |
| All | 381 | 1385 | 3182 | 4948 |

Of note, three B//LB variants were classified as P/LP by VIP-HL. The first one was NM_153676.3:c.2547-1G>T in the *USH1C* gene. It was submitted as likely benign in ClinVar, whereas VIP-HL assigned PVS1 and PM2, leading to a likely pathogenic classification. Manual curation revealed that this splicing variant affected an exon in transcript NM_153676.3 with no detectable expression based on the Genotype-Tissue Expression (GTEx) database ^32^ Thus, this variant should be classified as benign/likely benign. This example highlights the need for including information about the most biologically relevant transcripts in VIP-HL for accurate clinical variant interpretation^33^.

The other two discrepant variants were synonymous (NM_206933.3:c.949C>A (p.Arg317=) in the *USH2A* gene and NM_022124.6:c.7362G>A (p.Thr2454=) in the *CDH23* gene. Both were assigned likely benign classifications by VIP-HL due to the activation of BP4 and BP7. Specifically, NM_206933.3:c.949C>A was submitted as a pathogenic variant in ClinVar. This variant led to abnormal splicing and a premature termination codon,^34^ strongly supporting its pathogenic classification. As shown below, our VIP-HL user interface will enable curators to manually activate codes for functional studies to avoid such possible misclassifications. The second variant, NM_022124.6:c.7362G>A, was submitted as likely pathogenic No supporting RNA/functional, case-level or segregation studies were provided to support this classification. Since the actual effect of this sequence change is still unknown, a likely benign classification may be more appropriate for this case.

Of 4948 ClinVar variants, the most utilized rules were population-related, spanning PM2, BA1, BS1, BS2 and their modified strength levels (supporting through very strong; Figure 2). BP4 and BP7 are also frequently used because 33.5% (1657/4948) deafness-related variants with ClinVar star 2+ were synonymous variants. As expected, the pathogenic criteria were enriched in pathogenic/likely pathogenic variants, whereas benign criteria were enriched in benign/likely benign variants (Figure 2).

**Figure 2.**
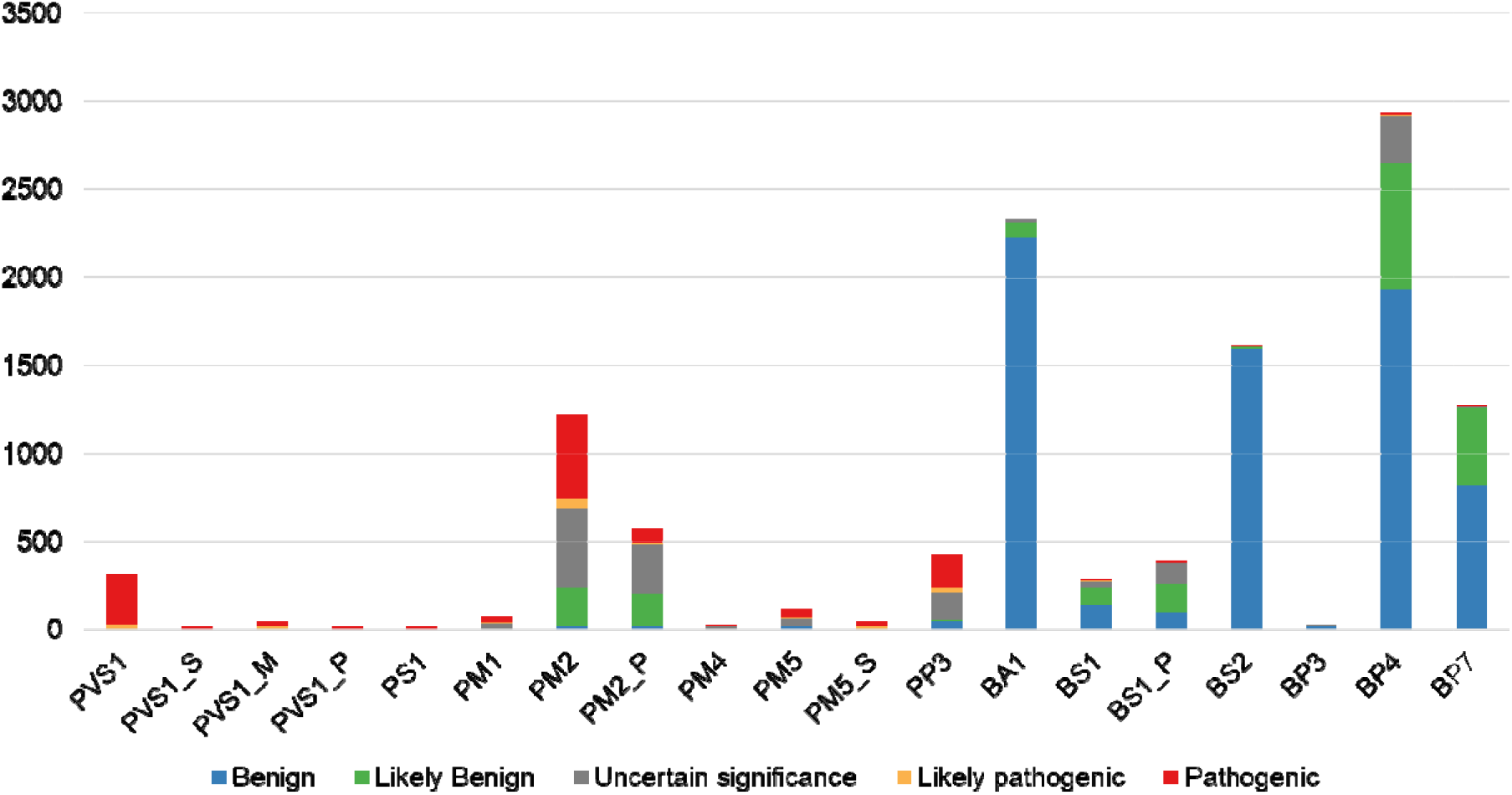
Frequency of rules applied for 4948 ClinVar variants. Rules applied with a modified strength are denoted by the rule followed by _P for Supporting, _M for Moderate, _S for Strong, and _VS for Very Strong.

### Features of VIP-HL web interface

Users can search the interface (http://hearing.genetics.bgi.com/) by gene name, chromosomal location, dbSNP identifier, and HGVS nomenclature.^35^ If a gene is queried, the results will present the gene name, cytobands, protein name, and external databases. All the pre-annotated variants are displayed with their pathogenicity and corresponding ACMG/AMP rules. If a specific variant is queried, VIP-HL displays the variant location (variant identifier), gene name, cHGVS, pHGVS, ACMG/AMP criteria, and classifications (Figure 3).

**Figure 3.**
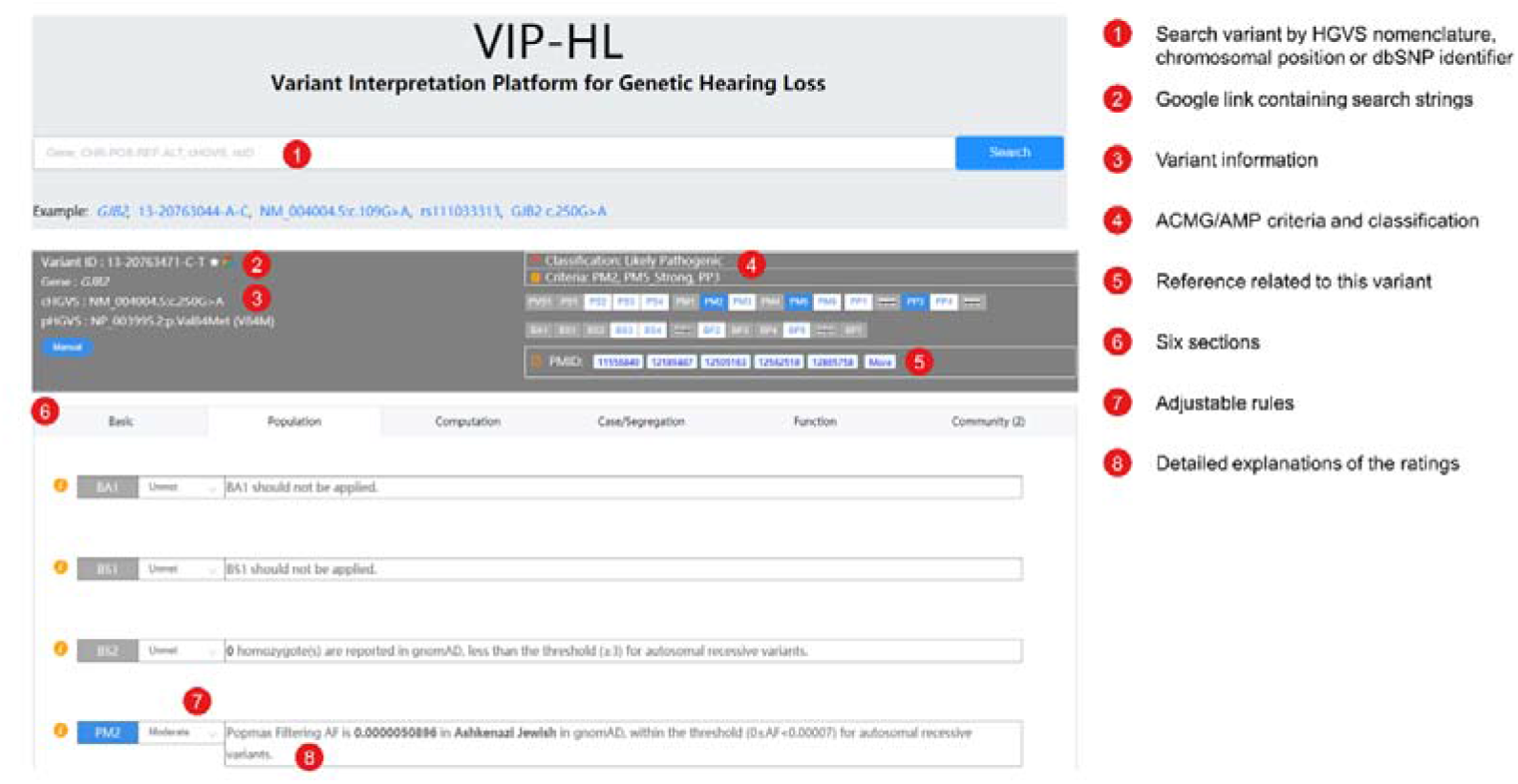
VIP-HL web-interface workflow. A variant could be searched by HGVS nomenclature, chromosomal location, or dbSNP identifier. Once a variant is queried, results are showed in three parts. The top-left corner displays variant information; the top-right corner displays ACMG/AMP criteria and final classification; the lower part displays six sections, including basic information, population, computation, case/segregation, function, and community. In each section, the strength of each criterion is adjustable based on users’ evaluations.

The browser displays six sections, including necessary information, population, computation, case/segregation, function, and community sections (Figure 3). The automatically annotated ACMG/AMP rules are presented in the population and computation sections, providing detailed explanations of the ratings based on the guidelines of genetic hearing loss.^7^ Rules that require manual curation are presented in the case/segregation and function sections. The community section records the submissions by users who would like to share their classifications with others.

Users are free to adjust the evidence strength from “Supporting” to “VeryStrong” in 24 ACMG/AMP rules. The classifications will update instantly with any rule(s) modifications. For rules grouped in the case/segregation and function sections, evidence to support the strength level can be manually added, enabling users to record and manage their curations.

## Discussion

Considering the substantial differences amongst diseases in terms of inheritance pattern, disease mechanism, phenotype, genetic and allelic heterogeneity, and prevalence, disease-specific guidelines are necessary for accurate and reliable interpretations.^36^ Following variant interpretation guidelines for genetic hearing loss,^7^ we developed a new computational tool, named VIP-HL, publicly available through a web interface (http://hearing.genetics.bgi.com/). To our knowledge, this is the first tool designed for automated variant interpretation in genetic hearing loss. Considering the high prevalence of hearing loss in the population, the availability of VIP-HL will significantly relieve the interpretation burdens for clinicians and curators.

Compared to rules activated by ClinGen HL-EP, VIP-HL showed a markedly high concordance (96%), indicating the reliability of interpreting hearing loss variants via VIP-HL. Of note, all the three discrepant activations (variants #1-3, Table 1) were attributable to population-based rules (BA1, and PM2), which depends on the adoption of popmax filtering allele frequency in extensive population studies.^18^ The ClinGen HL-EP used the ExAC database in the time of their research whereas we employed a larger dataset (gnomAD) as it was encouraged by the ClinGen HL-EP.^7^ Using these stringent allele frequencies empowers clinical genome interpretation without the removal of true pathogenic variants.^37^

VIP-HL activated several rules that were not activated by ClinGen HL-EP, including PM1 and BS2. ClinGen HL-EP did not perform a systematic review of mutational hot spots or functional domains for all genes associated with hearing loss, and proposed that PM1 can be applied for *KCNQ4* pore-forming region.^7^ In this study, we used the enrichment of pathogenic/likely pathogenic variants to construct a set of important regions ^19^ which includes the *KCNQ4* pore-forming region. Additionally, although HL-EP did not elaborate on the cutoff for BS2, we used a conservative cutoff to automate this rule. It should be noted that the penetrance affects the application of BS2 but was not considered by VIP-HL. This led to activations of BS2 for NM_004004.6:c.109G>A and NM_004004.6:c.101T>C in the *GJB2* gene because 50 and 16 homozygotes were identified from the gnomAD control dataset, respectively. The two variants were well-known pathogenic variants with low penetrance.^38^ Nevertheless, VIP-HL is a semi-automatic tool and our user interface enables curators to manually adjust codes to avoid such possible misclassifications.

A further comparison between VIP-HL and ClinVar showed an overall interpretation concordance of 88.0% based solely on the automation of only 13 ACMG/AMP rules. In terms of pathogenic/likely pathogenic variants, the concordance was much lower (57.1%). This could be explained by the lack of segregation and functional evidence from scientific literature, which requires manual curation and is a time-consuming process. Prospectively, text-mining and machine learning techniques might serve as potential solutions. For example, Birgmeier and co-authors developed an end-to-end machine learning tool, named AVADA, for the automatic retrieval of variant evidence directly from full-text literature.^39^ Suppose we can accumulate enormous datasets of evidence-related sentences or figures, in that case, it is possible to apply machine-learning approaches in the future for evidence retrieval and to automate the remaining ACMG/AMP rules in the next version of VIP-HL. In the meantime, our interface enables curators to manually activate the relevant codes after manual literature curation.

VIP-HL generated three P/LP classifications versus B/LB in ClinVar. All the three variants were related to the consideration of splicing impact. This discrepancy of NM_153676.3:c.2547-1G>T was attributable to a lack of considerations of exon expression data, which ultimately led to inappropriate classifications. It is apparent that a splicing variant affecting a non-expressive exon should have less functional effects.^33^ Recently, the transcript-level information from the GTEx project^32^ was utilized and proved that incorporating exon expression data can improve interpretations of putative loss-of-function variants.^40^ The second and third variants (NM_206933.3:c.949C>A and NM_022124.6:c.7362G>A) were synonymous variants, and their splicing impact should be curated from public literature if available. Nevertheless, these results indicated the importance of expression data in variant interpretation.

To improve user experience and further facilitate variation interpretation via VIP-HL, we developed a user-friendly web interface, which we continue to grow and add useful features over time. For example, PM3, one of the most frequently activated rules in genetic hearing loss,^7^ relies on the variant’s pathogenicity on the second allele. If this latter variant is introduced (in HGVS nomenclature) during the curation of PM3, VIP-HL can now provide the pathogenicity of this second variant as a reference for users. We expect such features and ongoing improvements would save curators the time and relieve the burden of variant interpretation.

VIP-HL has limitations. First, it is currently not applicable for exon-level copy number variations. Second, the allele frequency cutoffs were different for dominant and recessive hearing loss disorders. We first applied the cutoffs from the inheritance curated by ClinGen HL-EP for variants in a gene with both dominant and recessive inheritance. If both were available, we conservatively chose the cutoffs in recessive disorders. To avoid users falling into this pitfall, we highlighted the selected inheritance in the web interface of VIP-HL. Finally, the automation of rules relied on public databases. For example, PS1 and PM5 relied on ClinVar, which was recently reported to have misclassified variants,^28, 29^ which may lead to mis-annotations.

In conclusion, VIP-HL is an integrated online tool and search engine for variants in genetic hearing loss genes. It is also the first tool, to our knowledge, to consider the specifications proposed by ClinGen HL-EP for genetic hearing loss related variants. Providing reliable and reproducible annotations, VIP-HL not only facilitates variant interpretation but also provides a platform for users to share classifications with others.

