## Supplementary data for "VIP-HL: Semi-automated ACMG/AMP variant interpretation platform for genetic hearing loss"

Supplementary Figure 1. Frequency of rules exclusively activated by InterVar for 50 pilot variants

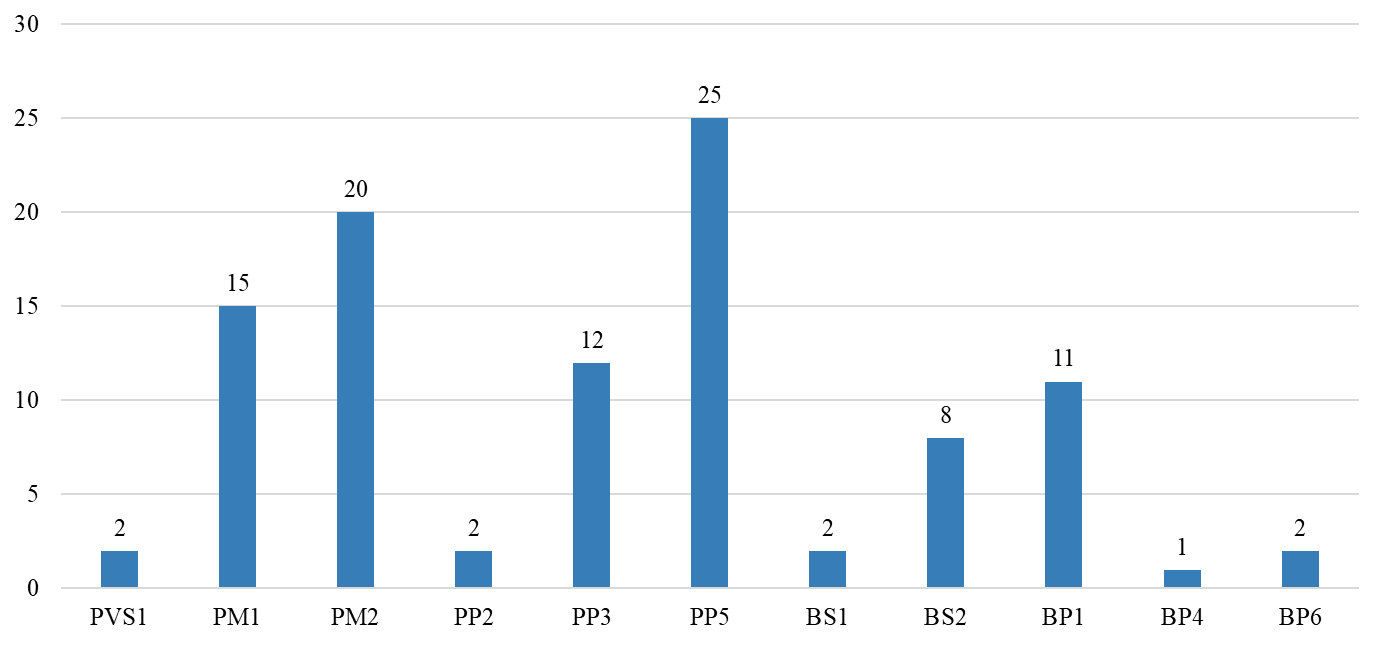

Supplementary Table 1. Gene-disease associations curated by ClinGen Hearing Loss Expert Panel

| # | Gene | Disease | Inheritance | Clinical validity | Classification date |
| --- | --- | --- | --- | --- | --- |
| 1 | [*ABHD12*](https://search.clinicalgenome.org/kb/genes/HGNC:15868) | PHARC syndrome | AR | Definitive | 2018-06-28 |
| 2 | [*ACTG1*](https://search.clinicalgenome.org/kb/genes/HGNC:144) | Nonsyndromic genetic deafness | AD | Definitive | 2019-01-07 |
| 3 | [*ACTG1*](https://search.clinicalgenome.org/kb/genes/HGNC:144) | Baraitser-winter syndrome 2 | AD | Definitive | 2019-01-07 |
| 4 | [*ADCY1*](https://search.clinicalgenome.org/kb/genes/HGNC:232) | Autosomal recessive nonsyndromic deafness | AR | Limited | 2017-05-10 |
| 5 | [*ADGRV1*](https://search.clinicalgenome.org/kb/genes/HGNC:17416) | Usher syndrome type 2 | AR | Definitive | 2017-02-15 |
| 6 | [*AIFM1*](https://search.clinicalgenome.org/kb/genes/HGNC:8768) | X-linked hereditary sensory and autonomic neuropathy with deafness | XL | Definitive | 2018-07-09 |
| 7 | [*ATP6V1B1*](https://search.clinicalgenome.org/kb/genes/HGNC:853) | Renal tubular acidosis, distal, with progressive nerve deafness | AR | Definitive | 2017-12-19 |
| 8 | [*BCS1L*](https://search.clinicalgenome.org/kb/genes/HGNC:1020) | Bjornstad syndrome | AR | Definitive | 2018-07-09 |
| 9 | [*BDP1*](https://search.clinicalgenome.org/kb/genes/HGNC:13652) | Nonsyndromic genetic deafness | AR | Limited | 2017-09-11 |
| 10 | [*BSND*](https://search.clinicalgenome.org/kb/genes/HGNC:16512) | Bartter disease type 4a | AR | Definitive | 2018-07-12 |
| 11 | [*BTD*](https://search.clinicalgenome.org/kb/genes/HGNC:1122) | Biotinidase deficiency | AR | Definitive | 2020-02-10 |
| 12 | [*CABP2*](https://search.clinicalgenome.org/kb/genes/HGNC:1385) | Nonsyndromic genetic deafness | AR | Definitive | 2020-02-06 |
| 13 | [*CACNA1D*](https://search.clinicalgenome.org/kb/genes/HGNC:1391) | Sinoatrial node dysfunction and deafness | AR | Moderate | 2018-05-01 |
| 14 | [*CCDC50*](https://search.clinicalgenome.org/kb/genes/HGNC:18111) | Nonsyndromic genetic deafness | AD | Limited | 2017-12-19 |
| 15 | [*CD164*](https://search.clinicalgenome.org/kb/genes/HGNC:1632) | Nonsyndromic genetic deafness | AD | Limited | 2018-03-20 |
| 16 | [*CDC14A*](https://search.clinicalgenome.org/kb/genes/HGNC:1718) | Nonsyndromic genetic deafness | AR | Limited | 2018-02-26 |
| 17 | [*CDC14A*](https://search.clinicalgenome.org/kb/genes/HGNC:1718) | Hearing impairment and infertile male syndrome | AR | Strong | 2018-02-26 |
| 18 | [*CDH23*](https://search.clinicalgenome.org/kb/genes/HGNC:13733) | Usher syndrome type 1 | AR | Definitive | 2017-01-30 |
| 19 | [*CDH23*](https://search.clinicalgenome.org/kb/genes/HGNC:13733) | Nonsyndromic genetic deafness | AR | Definitive | 2018-05-22 |
| 20 | [*CEACAM16*](https://search.clinicalgenome.org/kb/genes/HGNC:31948) | Nonsyndromic genetic deafness | AD | Moderate | 2018-03-20 |
| 21 | [*CEP78*](https://search.clinicalgenome.org/kb/genes/HGNC:25740) | Cone-rod dystrophy and hearing loss; CRDHL | AR | Strong | 2017-04-12 |
| 22 | [*CHD7*](https://search.clinicalgenome.org/kb/genes/HGNC:20626) | CHARGE syndrome | AD | Definitive | 2018-08-22 |
| 23 | [*CIB2*](https://search.clinicalgenome.org/kb/genes/HGNC:24579) | Nonsyndromic genetic deafness | AR | Definitive | 2018-02-20 |
| 24 | [*CISD2*](https://search.clinicalgenome.org/kb/genes/HGNC:24212) | Wolfram syndrome | AR | Strong | 2018-03-27 |
| 25 | [*CLDN14*](https://search.clinicalgenome.org/kb/genes/HGNC:2035) | Nonsyndromic genetic deafness | AR | Definitive | 2018-05-01 |
| 26 | [*CLIC5*](https://search.clinicalgenome.org/kb/genes/HGNC:13517) | Autosomal recessive nonsyndromic deafness | AR | Moderate | 2017-11-21 |
| 27 | [*CLPP*](https://search.clinicalgenome.org/kb/genes/HGNC:2084) | Perrault syndrome 3 | AR | Definitive | 2018-03-27 |
| 28 | [*CLRN1*](https://search.clinicalgenome.org/kb/genes/HGNC:12605) | Usher syndrome type 3 | AR | Definitive | 2017-03-02 |
| 29 | [*COCH*](https://search.clinicalgenome.org/kb/genes/HGNC:2180) | Nonsyndromic genetic deafness | AD | Definitive | 2018-01-05 |
| 30 | [*COL11A2*](https://search.clinicalgenome.org/kb/genes/HGNC:2187) | Nonsyndromic genetic deafness | AR | Moderate | 2018-12-20 |
| 31 | [*COL11A2*](https://search.clinicalgenome.org/kb/genes/HGNC:2187) | Nonsyndromic genetic deafness | AD | Moderate | 2018-12-20 |
| 32 | [*COL11A2*](https://search.clinicalgenome.org/kb/genes/HGNC:2187) | Otospondylomegaepiphyseal dysplasia | AR | Definitive | 2018-12-20 |
| 33 | [*COL11A2*](https://search.clinicalgenome.org/kb/genes/HGNC:2187) | Otospondylomegaepiphyseal dysplasia | AD | Definitive | 2018-12-20 |
| 34 | [*COL4A5*](https://search.clinicalgenome.org/kb/genes/HGNC:2207) | Alport syndrome | XL | Definitive | 2019-03-19 |
| 35 | [*COL4A6*](https://search.clinicalgenome.org/kb/genes/HGNC:2208) | Deafness, X-linked 6 | XL | Limited | 2018-01-16 |
| 36 | [*COL9A2*](https://search.clinicalgenome.org/kb/genes/HGNC:2218) | Stickler syndrome | AR | Limited | 2019-02-19 |
| 37 | [*COL9A3*](https://search.clinicalgenome.org/kb/genes/HGNC:2219) | Stickler syndrome | AR | Moderate | 2019-09-17 |
| 38 | [*CRYM*](https://search.clinicalgenome.org/kb/genes/HGNC:2418) | Autosomal dominant nonsyndromic deafness 40 | AD | Limited | 2017-02-16 |
| 39 | [*DCDC2*](https://search.clinicalgenome.org/kb/genes/HGNC:18141) | Nonsyndromic genetic deafness | AR | Limited | 2017-11-21 |
| 40 | [*DIABLO*](https://search.clinicalgenome.org/kb/genes/HGNC:21528) | Nonsyndromic genetic deafness | AD | Limited | 2017-12-19 |
| 41 | [*DIAPH1*](https://search.clinicalgenome.org/kb/genes/HGNC:2876) | Diaph1-related sensorineural hearing loss-thrombocytopenia syndrome | AD | Definitive | 2018-01-05 |
| 42 | [*DIAPH3*](https://search.clinicalgenome.org/kb/genes/HGNC:15480) | Auditory neuropathy | AD | Limited | 2020-02-06 |
| 43 | [*DMXL2*](https://search.clinicalgenome.org/kb/genes/HGNC:2938) | Nonsyndromic genetic deafness | AD | Limited | 2020-02-06 |
| 44 | [*DNMT1*](https://search.clinicalgenome.org/kb/genes/HGNC:2976) | Autosomal dominant cerebellar ataxia, deafness and narcolepsy | AD | Definitive | 2017-02-10 |
| 45 | [*DSPP*](https://search.clinicalgenome.org/kb/genes/HGNC:3054) | Dentinogenesis imperfecta (disease) | AD | Definitive | 2018-04-17 |
| 46 | [*EDN3*](https://search.clinicalgenome.org/kb/genes/HGNC:3178) | Waardenburg syndrome type 4B | AD | Limited | 2018-05-30 |
| 47 | [*EDN3*](https://search.clinicalgenome.org/kb/genes/HGNC:3178) | Waardenburg syndrome type 4B | AR | Moderate | 2018-05-08 |
| 48 | [*EDNRB*](https://search.clinicalgenome.org/kb/genes/HGNC:3180) | Waardenburg syndrome type 4A | AR | Moderate | 2018-05-08 |
| 49 | [*EDNRB*](https://search.clinicalgenome.org/kb/genes/HGNC:3180) | Waardenburg syndrome type 4A | AD | Limited | 2018-05-08 |
| 50 | [*ELMOD3*](https://search.clinicalgenome.org/kb/genes/HGNC:26158) | Nonsyndromic genetic deafness | AR | Limited | 2017-05-04 |
| 51 | [*EPS8*](https://search.clinicalgenome.org/kb/genes/HGNC:3420) | Autosomal recessive nonsyndromic deafness 102 | AR | Moderate | 2020-02-06 |
| 52 | [*EPS8L2*](https://search.clinicalgenome.org/kb/genes/HGNC:21296) | Nonsyndromic genetic deafness | AR | Moderate | 2020-02-05 |
| 53 | [*ESPN*](https://search.clinicalgenome.org/kb/genes/HGNC:13281) | Nonsyndromic genetic deafness | AD | Limited | 2018-09-20 |
| 54 | [*ESPN*](https://search.clinicalgenome.org/kb/genes/HGNC:13281) | Nonsyndromic genetic deafness | AR | Definitive | 2018-02-27 |
| 55 | [*ESRRB*](https://search.clinicalgenome.org/kb/genes/HGNC:3473) | Nonsyndromic genetic deafness | AR | Definitive | 2018-04-24 |
| 56 | [*EYA1*](https://search.clinicalgenome.org/kb/genes/HGNC:3519) | Branchio-oto-renal syndrome | AD | Definitive | 2018-08-30 |
| 57 | [*EYA4*](https://search.clinicalgenome.org/kb/genes/HGNC:3522) | Nonsyndromic genetic deafness | AD | Definitive | 2018-01-05 |
| 58 | [*FGF3*](https://search.clinicalgenome.org/kb/genes/HGNC:3681) | Deafness with labyrinthine aplasia, microtia, and microdontia | AR | Definitive | 2019-05-21 |
| 59 | [*FOXI1*](https://search.clinicalgenome.org/kb/genes/HGNC:3815) | Syndromic genetic deafness | AR | Limited | 2018-02-27 |
| 60 | [*GATA3*](https://search.clinicalgenome.org/kb/genes/HGNC:4172) | Hypoparathyroidism-deafness-renal disease syndrome | AD | Definitive | 2019-06-18 |
| 61 | [*GIPC3*](https://search.clinicalgenome.org/kb/genes/HGNC:18183) | Nonsyndromic genetic deafness | AR | Definitive | 2017-08-22 |
| 62 | [*GJB2*](https://search.clinicalgenome.org/kb/genes/HGNC:4284) | Autosomal recessive nonsyndromic deafness | AR | Definitive | 2017-03-02 |
| 63 | [*GJB2*](https://search.clinicalgenome.org/kb/genes/HGNC:4284) | Syndromic genetic deafness | AD | Definitive | 2018-06-26 |
| 64 | [*GPSM2*](https://search.clinicalgenome.org/kb/genes/HGNC:29501) | Chudley-mccullough syndrome | AR | Definitive | 2018-05-01 |
| 65 | [*GRHL2*](https://search.clinicalgenome.org/kb/genes/HGNC:2799) | Nonsyndromic genetic deafness | AD | Strong | 2018-01-16 |
| 66 | [*GRXCR1*](https://search.clinicalgenome.org/kb/genes/HGNC:31673) | Nonsyndromic genetic deafness | AR | Definitive | 2018-04-24 |
| 67 | [*GRXCR2*](https://search.clinicalgenome.org/kb/genes/HGNC:33862) | Nonsyndromic genetic deafness | AR | Moderate | 2019-01-07 |
| 68 | [*GSDME*](https://search.clinicalgenome.org/kb/genes/HGNC:2810) | Autosomal dominant nonsyndromic deafness | AD | Definitive | 2018-07-19 |
| 69 | [*HARS2*](https://search.clinicalgenome.org/kb/genes/HGNC:4817) | Perrault syndrome 2 | AR | Limited | 2018-05-15 |
| 70 | [*HGF*](https://search.clinicalgenome.org/kb/genes/HGNC:4893) | Nonsyndromic genetic deafness | AR | Moderate | 2018-01-16 |
| 71 | [*HOMER2*](https://search.clinicalgenome.org/kb/genes/HGNC:17513) | Nonsyndromic genetic deafness | AD | Moderate | 2018-10-16 |
| 72 | [*HSD17B4*](https://search.clinicalgenome.org/kb/genes/HGNC:5213) | Perrault syndrome | AR | Definitive | 2018-05-30 |
| 73 | [*ILDR1*](https://search.clinicalgenome.org/kb/genes/HGNC:28741) | Nonsyndromic genetic deafness | AR | Definitive | 2017-11-21 |
| 74 | [*KARS*](https://search.clinicalgenome.org/kb/genes/#N/A) | Nonsyndromic genetic deafness | AR | Limited | 2018-08-24 |
| 75 | [*KCNE1*](https://search.clinicalgenome.org/kb/genes/HGNC:6240) | Jervell and Lange-Nielsen syndrome 2 | AR | Moderate | 2018-06-22 |
| 76 | [*KCNQ1*](https://search.clinicalgenome.org/kb/genes/HGNC:6294) | Jervell and Lange-Nielsen syndrome | AR | Definitive | 2017-12-19 |
| 77 | [*KCNQ4*](https://search.clinicalgenome.org/kb/genes/HGNC:6298) | Nonsyndromic genetic deafness | AD | Definitive | 2017-11-21 |
| 78 | [*KITLG*](https://search.clinicalgenome.org/kb/genes/HGNC:6343) | Nonsyndromic genetic deafness | AD | Limited | 2018-01-16 |
| 79 | [*LARS2*](https://search.clinicalgenome.org/kb/genes/HGNC:17095) | Perrault syndrome | AR | Strong | 2018-06-27 |
| 80 | [*LHFPL5*](https://search.clinicalgenome.org/kb/genes/HGNC:21253) | Nonsyndromic genetic deafness | AR | Definitive | 2018-04-24 |
| 81 | [*LOXHD1*](https://search.clinicalgenome.org/kb/genes/HGNC:26521) | Nonsyndromic genetic deafness | AR | Definitive | 2018-05-08 |
| 82 | [*LRTOMT*](https://search.clinicalgenome.org/kb/genes/HGNC:25033) | Autosomal recessive nonsyndromic deafness 63 | AR | Definitive | 2017-02-15 |
| 83 | [*MARVELD2*](https://search.clinicalgenome.org/kb/genes/HGNC:26401) | Nonsyndromic genetic deafness | AR | Definitive | 2018-08-29 |
| 84 | [*MCM2*](https://search.clinicalgenome.org/kb/genes/HGNC:6944) | Autosomal dominant nonsyndromic deafness 70 | AD | Limited | 2018-01-04 |
| 85 | [*MET*](https://search.clinicalgenome.org/kb/genes/HGNC:7029) | Nonsyndromic genetic deafness | AR | Limited | 2017-12-19 |
| 86 | [*MITF*](https://search.clinicalgenome.org/kb/genes/HGNC:7105) | Waardenburg syndrome type 2 | AD | Definitive | 2018-07-26 |
| 87 | [*MSRB3*](https://search.clinicalgenome.org/kb/genes/HGNC:27375) | Nonsyndromic genetic deafness | AR | Moderate | 2017-11-15 |
| 88 | [*MYH14*](https://search.clinicalgenome.org/kb/genes/HGNC:23212) | Nonsyndromic genetic deafness | AD | Definitive | 2018-06-19 |
| 89 | [*MYH9*](https://search.clinicalgenome.org/kb/genes/HGNC:7579) | MYH-9 related disease | AD | Definitive | 2018-07-17 |
| 90 | [*MYO15A*](https://search.clinicalgenome.org/kb/genes/HGNC:7594) | Nonsyndromic genetic deafness | AR | Definitive | 2017-11-21 |
| 91 | [*MYO3A*](https://search.clinicalgenome.org/kb/genes/HGNC:7601) | Nonsyndromic genetic deafness | AR | Strong | 2017-11-15 |
| 92 | [*MYO6*](https://search.clinicalgenome.org/kb/genes/HGNC:7605) | Nonsyndromic genetic deafness | AD | Definitive | 2018-02-20 |
| 93 | [*MYO7A*](https://search.clinicalgenome.org/kb/genes/HGNC:7606) | Usher syndrome type 1 | AR | Definitive | 2018-06-29 |
| 94 | [*MYO7A*](https://search.clinicalgenome.org/kb/genes/HGNC:7606) | Nonsyndromic genetic deafness | AD | Definitive | 2018-03-19 |
| 95 | [*NARS2*](https://search.clinicalgenome.org/kb/genes/HGNC:26274) | Nonsyndromic genetic deafness | AR | Limited | 2017-12-19 |
| 96 | [*OSBPL2*](https://search.clinicalgenome.org/kb/genes/HGNC:15761) | Nonsyndromic genetic deafness | AD | Moderate | 2020-02-06 |
| 97 | [*OTOA*](https://search.clinicalgenome.org/kb/genes/HGNC:16378) | Nonsyndromic genetic deafness | AR | Definitive | 2018-05-01 |
| 98 | [*OTOF*](https://search.clinicalgenome.org/kb/genes/HGNC:8515) | Autosomal recessive nonsyndromic deafness 9 | AR | Definitive | 2017-01-30 |
| 99 | [*OTOG*](https://search.clinicalgenome.org/kb/genes/HGNC:8516) | Nonsyndromic genetic deafness | AR | Definitive | 2018-06-28 |
| 100 | [*OTOGL*](https://search.clinicalgenome.org/kb/genes/HGNC:26901) | Nonsyndromic genetic deafness | AR | Definitive | 2018-01-05 |
| 101 | [*P2RX2*](https://search.clinicalgenome.org/kb/genes/HGNC:15459) | Nonsyndromic genetic deafness | AD | Moderate | 2018-02-20 |
| 102 | [*PAX3*](https://search.clinicalgenome.org/kb/genes/HGNC:8617) | Waardenburg syndrome | AD | Definitive | 2017-11-15 |
| 103 | [*PCDH15*](https://search.clinicalgenome.org/kb/genes/HGNC:14674) | Usher syndrome type 1 | AR | Definitive | 2017-02-15 |
| 104 | [*PCDH15*](https://search.clinicalgenome.org/kb/genes/HGNC:14674) | Nonsyndromic genetic deafness | AR | Limited | 2018-06-19 |
| 105 | [*PDZD7*](https://search.clinicalgenome.org/kb/genes/HGNC:26257) | Autosomal recessive nonsyndromic deafness | AR | Definitive | 2017-04-26 |
| 106 | [*PJVK*](https://search.clinicalgenome.org/kb/genes/HGNC:29502) | Nonsyndromic genetic deafness | AR | Definitive | 2017-12-19 |
| 107 | [*PNPT1*](https://search.clinicalgenome.org/kb/genes/HGNC:23166) | Autosomal recessive nonsyndromic deafness | AR | Limited | 2017-02-23 |
| 108 | [*POU3F4*](https://search.clinicalgenome.org/kb/genes/HGNC:9217) | Nonsyndromic genetic deafness | XL | Definitive | 2018-01-05 |
| 109 | [*POU4F3*](https://search.clinicalgenome.org/kb/genes/HGNC:9220) | Nonsyndromic genetic deafness | AD | Definitive | 2017-11-21 |
| 110 | [*PRPS1*](https://search.clinicalgenome.org/kb/genes/HGNC:9462) | PRPS1 deficiency disorder | XL | Definitive | 2020-02-14 |
| 111 | [*PRPS1*](https://search.clinicalgenome.org/kb/genes/HGNC:9462) | Phosphoribosylpyrophosphate synthetase superactivity | XL | Limited | 2020-02-14 |
| 112 | [*PTPRQ*](https://search.clinicalgenome.org/kb/genes/HGNC:9679) | Autosomal recessive nonsyndromic deafness | AR | Definitive | 2017-02-23 |
| 113 | [*RDX*](https://search.clinicalgenome.org/kb/genes/HGNC:9944) | Nonsyndromic genetic deafness | AR | Definitive | 2018-01-02 |
| 114 | [*RIPOR2*](https://search.clinicalgenome.org/kb/genes/HGNC:13872) | Nonsyndromic genetic deafness | AR | Moderate | 2018-09-18 |
| 115 | [*ROR1*](https://search.clinicalgenome.org/kb/genes/HGNC:10256) | Nonsyndromic genetic deafness | AR | Limited | 2018-03-20 |
| 116 | [*S1PR2*](https://search.clinicalgenome.org/kb/genes/HGNC:3169) | Nonsyndromic genetic deafness | AR | Strong | 2017-03-02 |
| 117 | [*SERPINB6*](https://search.clinicalgenome.org/kb/genes/HGNC:8950) | Nonsyndromic genetic deafness | AR | Moderate | 2018-04-24 |
| 118 | [*SIX1*](https://search.clinicalgenome.org/kb/genes/HGNC:10887) | Branchio-oto-renal syndrome | AD | Definitive | 2017-06-22 |
| 119 | [*SLC17A8*](https://search.clinicalgenome.org/kb/genes/HGNC:20151) | Nonsyndromic genetic deafness | AD | Strong | 2018-09-18 |
| 120 | [*SLC26A4*](https://search.clinicalgenome.org/kb/genes/HGNC:8818) | Pendred syndrome | AR | Definitive | 2018-08-02 |
| 121 | [*SLC26A5*](https://search.clinicalgenome.org/kb/genes/HGNC:9359) | Nonsyndromic genetic deafness | AR | Limited | 2017-12-19 |
| 122 | [*SLC44A4*](https://search.clinicalgenome.org/kb/genes/HGNC:13941) | Nonsyndromic genetic deafness | AD | Limited | 2018-04-24 |
| 123 | [*SLC52A2*](https://search.clinicalgenome.org/kb/genes/HGNC:30224) | Brown-Vialetto-van Laere syndrome 2 | AR | Definitive | 2018-07-09 |
| 124 | [*SLC52A3*](https://search.clinicalgenome.org/kb/genes/HGNC:16187) | Brown-Vialetto-van Laere syndrome | AR | Definitive | 2018-09-18 |
| 125 | [*SLITRK6*](https://search.clinicalgenome.org/kb/genes/HGNC:23503) | High myopia-sensorineural deafness syndrome | AR | Strong | 2018-03-20 |
| 126 | [*SMPX*](https://search.clinicalgenome.org/kb/genes/HGNC:11122) | Nonsyndromic genetic deafness | XL | Definitive | 2017-09-12 |
| 127 | [*SNAI2*](https://search.clinicalgenome.org/kb/genes/HGNC:11094) | Waardenburg syndrome | AR | Limited | 2018-08-23 |
| 128 | [*SOX10*](https://search.clinicalgenome.org/kb/genes/HGNC:11190) | Waardenburg syndrome type 4C | AD | Definitive | 2018-06-19 |
| 129 | [*STRC*](https://search.clinicalgenome.org/kb/genes/HGNC:16035) | Nonsyndromic genetic deafness | AR | Definitive | 2017-12-19 |
| 130 | [*SYNE4*](https://search.clinicalgenome.org/kb/genes/HGNC:26703) | Nonsyndromic genetic deafness | AR | Moderate | 2020-02-06 |
| 131 | [*TBC1D24*](https://search.clinicalgenome.org/kb/genes/HGNC:29203) | Nonsyndromic genetic deafness | AD | Limited | 2018-03-27 |
| 132 | [*TCOF1*](https://search.clinicalgenome.org/kb/genes/HGNC:11654) | Treacher-Collins syndrome | AD | Definitive | 2019-09-17 |
| 133 | [*TECTA*](https://search.clinicalgenome.org/kb/genes/HGNC:11720) | Nonsyndromic genetic deafness | AR | Definitive | 2018-09-12 |
| 134 | [*TECTA*](https://search.clinicalgenome.org/kb/genes/HGNC:11720) | Nonsyndromic genetic deafness | AD | Definitive | 2018-01-02 |
| 135 | [*TIMM8A*](https://search.clinicalgenome.org/kb/genes/HGNC:11817) | Deafness dystonia syndrome | XL | Definitive | 2017-12-19 |
| 136 | [*TJP2*](https://search.clinicalgenome.org/kb/genes/HGNC:11828) | Nonsyndromic genetic deafness | AD | Limited | 2017-12-19 |
| 137 | [*TMC1*](https://search.clinicalgenome.org/kb/genes/HGNC:16513) | Autosomal recessive nonsyndromic deafness 7 | AR | Definitive | 2017-02-15 |
| 138 | [*TMC1*](https://search.clinicalgenome.org/kb/genes/HGNC:16513) | Nonsyndromic genetic deafness | AD | Definitive | 2018-06-22 |
| 139 | [*TMEM132E*](https://search.clinicalgenome.org/kb/genes/HGNC:26991) | Autosomal recessive nonsyndromic deafness | AR | Limited | 2017-11-21 |
| 140 | [*TMIE*](https://search.clinicalgenome.org/kb/genes/HGNC:30800) | Nonsyndromic genetic deafness | AR | Definitive | 2017-09-29 |
| 141 | [*TMPRSS3*](https://search.clinicalgenome.org/kb/genes/HGNC:11877) | Nonsyndromic genetic deafness | AR | Definitive | 2017-08-22 |
| 142 | [*TNC*](https://search.clinicalgenome.org/kb/genes/HGNC:5318) | Nonsyndromic genetic deafness | AD | Limited | 2018-03-20 |
| 143 | [*TPRN*](https://search.clinicalgenome.org/kb/genes/HGNC:26894) | Nonsyndromic genetic deafness | AR | Definitive | 2017-09-12 |
| 144 | [*TRIOBP*](https://search.clinicalgenome.org/kb/genes/HGNC:17009) | Autosomal recessive nonsyndromic deafness | AR | Definitive | 2017-06-06 |
| 145 | [*TUBB4B*](https://search.clinicalgenome.org/kb/genes/HGNC:20771) | Leber congenital amaurosis with early-onset deafness | AD | Moderate | 2018-06-26 |
| 146 | [*USH1C*](https://search.clinicalgenome.org/kb/genes/HGNC:12597) | Usher syndrome type 1 | AR | Definitive | 2017-02-15 |
| 147 | [*USH1C*](https://search.clinicalgenome.org/kb/genes/HGNC:12597) | Nonsyndromic genetic deafness | AR | Limited | 2018-06-11 |
| 148 | [*USH1G*](https://search.clinicalgenome.org/kb/genes/HGNC:16356) | Usher syndrome type 1 | AR | Definitive | 2017-02-15 |
| 149 | [*USH2A*](https://search.clinicalgenome.org/kb/genes/HGNC:12601) | Usher syndrome type 2 | AR | Definitive | 2017-02-15 |
| 150 | [*WBP2*](https://search.clinicalgenome.org/kb/genes/HGNC:12738) | Autosomal recessive nonsyndromic deafness | AR | Limited | 2018-12-09 |
| 151 | [*WFS1*](https://search.clinicalgenome.org/kb/genes/HGNC:12762) | Wolfram syndrome | AR | Definitive | 2018-09-11 |
| 152 | [*WFS1*](https://search.clinicalgenome.org/kb/genes/HGNC:12762) | Wolfram-like syndrome | AD | Definitive | 2018-04-17 |
| 153 | [*WFS1*](https://search.clinicalgenome.org/kb/genes/HGNC:12762) | Neonatal-onset diabetes, congenital sensorineural deafness, and congenital cataracts | AD | Strong | 2018-04-17 |
| 154 | [*WHRN*](https://search.clinicalgenome.org/kb/genes/HGNC:16361) | Usher syndrome type 2D | AR | Definitive | 2017-05-10 |
| 155 | [*WHRN*](https://search.clinicalgenome.org/kb/genes/HGNC:16361) | Nonsyndromic genetic deafness | AR | Moderate | 2018-12-26 |
| 156 | [*ALMS1*](https://search.clinicalgenome.org/kb/genes/HGNC:428) | Alstrom syndrome | AR | Definitive | 2017-02-10 |
| 157 | [*COL2A1*](https://search.clinicalgenome.org/kb/genes/HGNC:2200) | Spondyloepiphyseal dysplasia, Stanescu type | AD | Moderate | 2016-12-01 |
| 158 | [*COL9A1*](https://search.clinicalgenome.org/kb/genes/HGNC:2217) | Stickler syndrome | AR | Limited | 2018-03-26 |
| 159 | [*FGFR3*](https://search.clinicalgenome.org/kb/genes/HGNC:3690) | Achondroplasia | AD | Definitive | 2016-12-01 |
| 160 | [*NDP*](https://search.clinicalgenome.org/kb/genes/HGNC:7678) | Norrie disease | XL | Definitive | 2018-03-21 |
| 161 | [*MIR96*](https://search.clinicalgenome.org/kb/genes/HGNC:31648) | Nonsyndromic hearing loss | AD | Moderate | 2018-05-22 |
